# SLO2.1/NALCN Functional Complex Activity in Mouse Myometrial Smooth Muscle Cells During Pregnancy

**DOI:** 10.1101/2024.05.29.596465

**Authors:** Juan J. Ferreira, Lindsey N. Kent, Ronald McCarthy, Alice Butler, Xiaofeng Ma, Nikita Peramsetty, Chinwendu Amazu, Alexander Zhang, Grace C. Whitter, Ethan Li, Sarah K. England, Celia M. Santi

## Abstract

At the end of pregnancy, the uterus transitions from a quiescent to a highly contractile state. This is partly due to depolarization of the resting membrane potential in uterine (myometrial) smooth muscle cells (MSMCs). In human MSMCs, the membrane potential is regulated by a functional complex between the sodium (Na^+^)-activated potassium (K^+^) channel SLO2.1 and the Na^+^ Leak Channel Non-Selective (NALCN). Na^+^ entering through NALCN activates SLO2.1, leading to K^+^ efflux, membrane hyperpolarization (cells become more negative inside), and reduced contractility. Decreased SLO2.1/NALCN activity results in reduced K^+^ efflux, leading to membrane depolarization, Ca^2+^ influx via voltage-dependent calcium channels, and increased MSMC contractility. However, all of these data are from MSMCs isolated from women at term, so the role of the SLO2.1/NALCN complex early in pregnancy was speculative. To address this question here, we examined the role of the SLO2.1/NALCN complex in regulating mouse MSMC membrane potential across pregnancy. We report that *Slo2.1* and *Nalcn* are more highly expressed in MSMCs from non-pregnant and early pregnant mice than in those from late-pregnant mice. Functional studies revealed that SLO2.1 channels mediate a significant portion of the K^+^ current in mouse MSMCs, particularly in cells from non-pregnant and early pregnant mice. Activation of SLO2.1 by Na^+^ influx through NALCN led to membrane hyperpolarization in MSMCs from early pregnancy but not in MSMCs from later pregnancy. Moreover, the NALCN/SLO2.1 complex regulates intracellular Ca^2+^ responses more in MSMCs from non-pregnant and early pregnancy mice than in MSMCs from late pregnancy. Together, these findings reveal that the SLO2.1/NALCN functional complex is conserved between mouse and humans and functions throughout pregnancy. This work could open avenues for targeted pharmacological interventions for pregnancy-related complications.

## 1. Introduction

During much of pregnancy, the uterus is quiescent. At term, the uterus becomes highly contractile to expel the fetus (Casteels & Kuriyama, 1965; Parkington et al., 1999). Uterine contractility is partially controlled by myometrial smooth muscle cell (MSMC) excitability, which is regulated by the MSMC resting membrane potential (Vm) (Ferreira et al., 2019, 2021; Parkington et al., 1999; Reinl et al., 2015a, 2018). The Vm is maintained through a balance between hyperpolarizing forces, primarily driven by outward potassium (K^+^) currents (Anwer et al., 1993; Knock et al., 1999), and depolarizing forces, largely mediated by inward sodium (Na^+^) leak currents. Changes in these ion fluxes modify the Vm and modulate calcium (Ca^2+^) influx through voltage-dependent Ca^2+^ channels (VDCCs) and other Ca^2+^ channels such as TRP C1, 3, and 6 (Babich et al., 2004; Dalrymple, 2002; Dalrymple et al., 2007; Ku et al., 2006; Lacampagne et al., 1994; Wang et al., 2020; Yang & Sachs, 1989).

MSMCs isolated from non-laboring women at term exhibit an Na^+^-regulated K^+^ current that is lost when the K^+^ channel SLO2.1 is knocked down. The current through SLO2.1 hyperpolarizes the MSMC Vm and dampens excitability (Ferreira et al., 2019). SLO2.1 is activated by Na^+^ influx through sodium leak channel nonselective (NALCN). This was demonstrated by showing that inhibition of NALCN with gadolinium (Gd^2+^) or replacing Na^+^ with Lithium (Li^+^), which can pass through NALCN but not activate SLO2.1, prevented Na^+^-activated K^+^ influx. Furthermore, NALCN and SLO2.1 are in proximity to one another in human MSMCs, and functional coupling of NALCN and SLO2.1 regulates contractility (Ferreira et al., 2021). These data support a model that, during pregnancy, Na^+^ influx through NALCN activates SLO2.1, leading to K^+^ influx, Vm hyperpolarization, and MSMC quiescence. However, all these experiments were done with MSMCs from women at term, so the functional interactions between SLO2.1 and NALCN earlier in pregnancy are unknown.

In this study, we investigated SLO2.1/NALCN complex dynamics in a mouse model in which we could compare various stages of pregnancy. We report differential expression of both SLO2.1 and NALCN channels in mouse MSMCs at various pregnancy stages. mRNA expression is lowest in late pregnancy, when MSMCs are most excitable. Furthermore, we report a reduction in SLO2.1 currents and SLO2.1/NALCN complex activity in mouse MSMCs in late pregnancy. Together, our data indicate that regulation of uterine excitability through the SLO2.1/NALCN functional complex is conserved across species and that this complex is important throughout different stages of pregnancy.

## Methods

### 2.1. Animals and tissue collection

Animal procedures followed NIH guidelines and were approved by the Institutional Animal Care and Use Committee at Washington University. C57BL/6 mice (Jackson Laboratories) were housed in 12-hour light/dark cycle with *ad libitum* access to food and water. Each female mouse aged 8-22 weeks was housed with a male aged 8-22 weeks and checked every morning. The day on which a copulatory plug was detected was recorded as pregnancy day 0.5. Uteri were collected from non-pregnant mice at different stages of estrus and from pregnant mice at gestational days 7.5, 10.5, 14.5, and 18.5.

### 2.2. Mouse myometrium digestion and cell culture

The uterus was dissected from euthanized mice and opened longitudinally, with the longitudinal smooth muscle layer facing up. Fetuses, placenta, and decidua lining were meticulously removed from the uterus. The uterine muscle was then thoroughly washed with Hanks’ Balanced Salt Solution (HBSS) or Dulbecco’s Phosphate-Buffered Saline (DPBS) (Ca²⁺ and Mg²⁺ free) at least twice to eliminate blood and debris. The cleaned uterine tissue was cut into squares of approximately 2 mm to 5 mm.

Smooth muscle cells were dissociated in DPBS or Dulbecco’s Modified Eagle Medium (DMEM) supplemented with 0.5% Bovine Serum Albumin (BSA) and 0.25 units/ml of Liberase TM (Roche, 165 µg in 7.5 ml). The tissue pieces were incubated at 37°C with constant shaking for 45-60 minutes. Then, the tissue pieces were transferred to a fresh solution devoid of the enzyme. Using fire-polished Pasteur pipettes of varying sizes, the tissue was mechanically disaggregated by pipetting up and down for several minutes, gradually reducing the size of the pipette openings as the tissue disintegrated into a cellular suspension.

The cell suspension was then filtered through a 70-µm strainer to remove any undigested tissue pieces and centrifuged at 860 relative centrifugal force (RCF) for 10 minutes. After discarding the supernatant, the cell pellet was resuspended in 1-2 ml of fresh 10% DMEM-F12 media supplemented with 100 units/ml penicillin and 100 µg/ml streptomycin (Complete Media). The resuspended cells were then seeded in a T75 cm² flask containing 10-20 ml of complete DMEM-F12 media and incubated at 37°C with 5% CO₂ for a maximum of 14-18 hours before being used for experimental procedures.

### 2.3. RNA isolation, reverse transcription, and real time PCR

To isolate total RNA from primary mouse MSMCs and hTERT-HM cells, the Qiagen RNeasy Kit (catalogue no. 74124; Qiagen, Valencia, CA) was used. Superscript III Reverse Transcriptase (catalogue no. 18080-044; Invitrogen, Carlsbad, CA) and random hexa-nucleotide primers (catalogue no. H0268-1UN; Sigma-Aldrich, St Louis, MO) were used to synthesize cDNA. PCR primers (Integrated DNA Technologies, Coralville, IA) were designed with Oligo 7 software (Molecular Biology Insights, Colorado Springs, CO). RNA was extracted from uterine tissue with Trizol and the aurum total RNA fatty and Bio-rads fibrous tissue module. iScript reverse transcription supermix was used to synthesize cDNA. Real-time PCR was performed with iQ-SYBR Green Supermix on a CFX96 Real-time System in the following conditions: 95°C for 5 minutes, then 50 cycles of 95°C for 10 seconds and 60°C for 30 seconds. Reactions were performed in duplicate, and data were normalized to Top1 and Sdha using the delta-delta CT method. Primer sequences: *Nalcn*, 5’TTTCCCCGCTGGCGCTCCTA3’ and 5’ACCAGCTGCCAACCACCAGC3’; *Top1*, 5’AAGATCGAGAACACCGGCATA3’ and 5’ CTTTTCCTCCTTTCGTCTTTCC3’; *Sdha*, 5’GGAACACTCCAAAAACAGACCT3’ and 5’CCACCACTGGGTATTGAGTAGAA3’ (Reinl et al., 2018); *Slo2.1*, 5’TCTTGAGAGCATGGGCTGTG3’ and 5’GGTGACTGCTGCCCTTCTTG3’.

### 2.4. Immunohistochemistry

Uteri were fixed in 10% buffered formalin, processed, and sectioned (4 µm). Slides were incubated in citric acid-based solution at ∼95°C for 30 min, blocked with Avidin/Biotin kit and 10% normal goat serum for 30 min, and incubated overnight at 4°C in primary antibodies (1:500 anti-NALCN, Alomone labs; 1:4000 anti-SLO2.1, Absolute Antibody; 1:50 alpha-Smooth muscle actin, Invitrogen). Slides were incubated in biotinylated secondary antibodies in 10% normal goat serum for 30 min at room temperature. Slides were washed three times with PBS between each step except after the block. Signal was detected with Vectastain Elite ABC and DAB substrate kits. Nuclei were counterstained with Gills Hematoxylin. Images were captured on a Leica TCS SPE microscope.

### 2.5. Electrophysiology

Cells were serum starved in DMEM for at least 2 hours before patch clamp experiments. Borosilicate glass pipettes (Warner Instruments, Hamden, CT) with 1.0–1.8 megaohm resistance were used for whole-cell recordings, and pipettes with 4–6 megaohm resistance were used for single-channel recordings. Whole-cell recordings of mouse MSMCs were performed in asymmetrical K^+^ solutions (unless specified in the figures) as follows (in mM): External solution 135 NaCl (replaced by 135 LiCl, in the Na-free or 0 mM Na^+^ solution), 5 KCl, 5 Hepes, 2 MgCl_2_; Internal solution, 140 KCl, 5 Hepes, 0.5 MgCl_2_, 5 ATPMg (0.6 mM free Mg^2+^) and 10 EGTA (0 Ca^2+^ solution), or 1 EGTA and 100 nM Ca^2+^ free (Ca^2+^-containing solution). The pH of all solutions was adjusted to 7.3-7.4. During experiments, cells were continuously perfused. Traces were acquired with Axopatch 200B (Molecular Devices, Sunnyvale, CA), digitized at 10 kHz for whole-cell recordings. Records were filtered at 2 kHz for macro whole-cell currents. Data were analyzed with pClamp version 10.6 (Molecular Devices) and SigmaPlot version 15 (Systat Software Inc., Chicago, IL).

### 2.6. *In situ* proximity ligation assay

Mouse MSMCs were cultured in 8-well chambered slides and serum-deprived in 0.5% FBS for 24 hours. Cells were then washed in ice-cold 1X PBS, fixed with 4% paraformaldehyde in PBS for 20 minutes at room temperature, washed 4 x 5-minutes in 1X PBS, permeabilized with 0.1% NP-40 for 5 minutes at room temperature, washed twice with PBS and once with 100 mM Glycine in PBS, and rinsed in milliQ water. Duolink i*n situ* proximity ligation assay (PLA) labeling was performed with NALCN (mouse monoclonal, 1:100, StressMarq) and SLO2.1 (rabbit polyclonal, 1:200, Alomone) antibodies. The manufacturer’s protocol was followed except that cells were stained with NucBlue Fixed Cell Stain Ready Probes (Invitrogen) for 5 minutes at room temperature before the final wash in wash buffer B. Slides were dried at room temperature in the dark, mounted in Vectashield, and stored at −20°C until analysis. Images were captured with a Leica AF 6000LX system with a Leica DMi8000 inverted microscope, a 63X objective (HC PL FluoTar L 63X/0.70 Dry), and an Andor-Zyla-VCS04494 camera at room temperature. Excitation wavelengths were 488 nm for PLA signals and 340 nm for NucBlue. Leica LasX software controlled the system and collected data. Acquisition parameters were: 2- and 1.5-seconds exposure time for 488 nm and 340 nm, respectively, no binning, 2048 x 2048 pixels resolution, and a voxel size of 0.103 mm. Image analysis was performed with LAS X, ImageJ, and SigmaPlot 15. Data are reported as the number of punctae per cell after background subtraction.

### 2.7. Determination of Vm by flow cytometry

Mouse MSMCs were centrifuged at 325 RCF for 5 min and resuspended in modified Ringer’s solution (in mM, 80 Choline Cl, 10 HEPES, 5 Glucose, 5 KCl, 2 CaCl_2_, pH 7.4). Before recording, 0.02 mg/ml Hoechst and 150-500 nM DiSC3(5) were added to 500 μl of cell suspension. Data were recorded as individual cellular events based on Side Scatter Area (SSC-A) and Forward Scatter Area (FSC-A), with fluorescence collected from 10,000-25,000 events per condition. Thresholds for FSC-A and SSC-A were set to exclude signals from cellular debris. Living cells were selected with the Pacific Blue filter (450 nm), and DiSC3(5)-positive cells were detected with the Allophycocyanine APC filter (670 nm).

To measure the effect of Na^+^ on Vm, 80 mM NaCl or Lithium (Li^+^) was added to the 500 μl suspensions. At the end of each experiment, 1 µM Valinomycin was added to establish a reference point for cross-experiment comparisons. Normalization was calculated as ((F_Valino_ - F_Ref_)/(F_Valino_ - F_Cation_))*100, where F_Ref_ is the median fluorescence of the population in the basal condition, F_Cation_ is after addition of the cation, and F_Valino_ is after addition of Valinomycin. Data were analyzed with FlowJo 10.6.1 software and reported as median values.

### 2.8. Lentivirus production and injection

*Scrambled*, *Slo2.1,* and *Nalcn* shRNAs were cloned into the pGFP-C-shLenti vector (OriGene) and co-transfected with pMD2.G and psPAX2 (Addgene, plasmid # 12259 and 12260) into HEK293T cells to produce virus. Media from transfected HEK293T cells was collected and concentrated with Lenti-X concentrator. Lentiviruses were injected into multiple sites along the uterine horn in non-pregnant mice, and MSMCs were isolated 7-10 days later. EGFP positivity was used to identify MSMCs expressing the shRNAs. The following shRNA sequences were used: *Slo2.1,* TCTAACTTGGCCTTTATGTTTCGACTGCC and AGAGCCATTCAGCGTACACAGTCTGCAAT; *Nalcn,* TGGTGACTGTGGATGTAATAGTTGCTGCC and AACAGGACACCTGCTGTCTCTTCAGAATC.

### 2.9. Calcium imaging

Cells were cultured on glass coverslips for 16-24 hours at 37 °C, incubated in 2 μM Fluo-4 AM and 0.05-0.1% Pluronic Acid F-127 in Opti-Mem for 60-90 min, then incubated in Ringer’s solution for 10 to 20 minutes. Solutions were applied with a perfusion system with an exchange time of approximately 1.5 seconds. Recordings began 2-5 minutes before addition of test solutions. Ionomycin (2-5 μM) was added at the end of recordings as a reference point for comparison across experiments. Calcium (Ca^2+^) signals were recorded using a Leica AF 6000LX system with a Leica DMi8000 inverted microscope, a 40X (HC PL FluoTar L 40X/0.70 Dry) or a 20X (N-Plan L 20X/0.35 Dry) air objective, and an Andor-Zyla-VCS04494 camera. Excitation and emission filters were 488/20 nm and 530/20 nm, respectively. Leica LasX2.0.014332 software-controlled image collection. Acquisition parameters: 120 ms exposure time, 2×2 binning, 512 x 512 pixels resolution, with a voxel size of 1.3 mm for the 20X objective. Images were captured every 10 seconds. Data were analyzed with LAS X, ImageJ, Clampfit 10 (Molecular Devices), and SigmaPlot 15 software. Changes in intracellular Ca^2+^ concentration are presented as (F/F_Iono_) after background subtraction. All imaging experiments were conducted at room temperature. Cells were considered responsive if they exhibited fluorescence changes of at least 5-10% of ionomycin responses after normalization.

### 2.10. Statistical Analysis

Statistical analyses were performed using SigmaPlot (version 15.0). For independent samples, an unpaired Student’s t-test was used, while a paired t-test was applied for case-control designs involving the same animals. Comparisons among multiple groups were conducted using one-way ANOVA or appropriate multiple comparison tests. Data are presented as mean ± standard deviation, with P < 0.05 considered statistically significant. Plotted values represent the mean of all technical replicates (1–3) performed within each experiment. In Figures 1, 3, 5, and 6, values represent the average per animal, while in Figures 2 and 4, they correspond to the average per cell tested. Since no a priori power analysis was conducted to determine the required sample sizes, findings from sample sizes below five should be considered preliminary, and P-values should be interpreted as descriptive.

**Figure 1.**
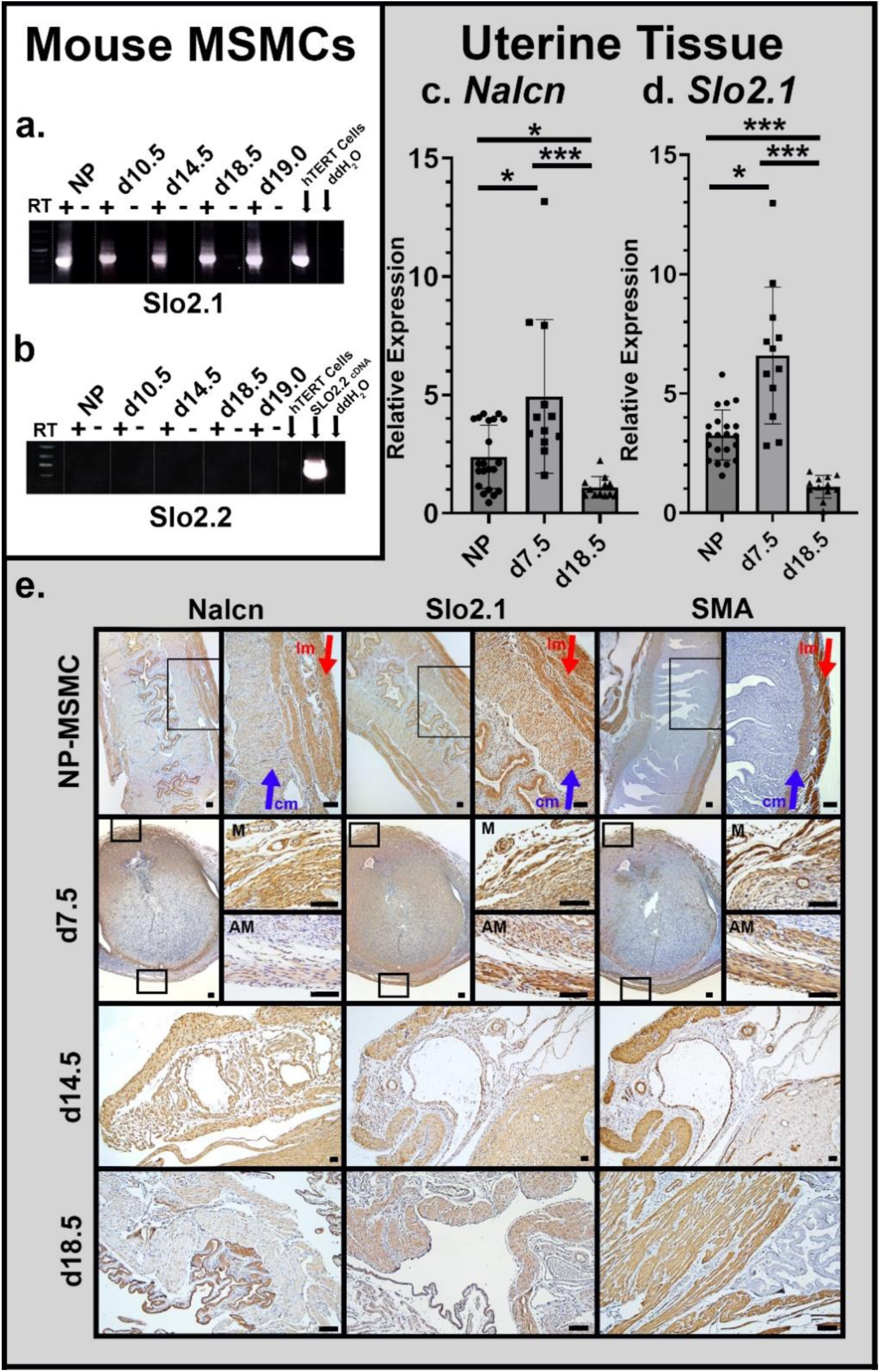
SLO2.1 and NALCN expression in uterus and myometrial smooth muscle cells. (**a** and **b**) Representative reverse transcription PCR of (**a**) *Slo2.1* or (**b**) *Slo2.2* in myometrial smooth muscle cells (MSMCs) from non-pregnant (NP) and day post coitus (d), 10.5, 14.5, 18.5, and 19 mice. No reverse transcriptase (RT) and ddH_2_O are negative controls, hTERT cell cDNA is a positive control. (**c** and **d**) Relative mRNA expression of (**c**) *Nalcn* and (**d**) *Slo2.1* in mouse uteri as measured by quantitative real-time PCR. Expression was normalized to *Top1* and *Sdha*. We analyzed NALCN and SLO2.1 mRNA expression in 21 non-pregnant (NP), 12 day 7.5 (d7.5), and 12 day 18.5 (d18.5) mice. \**P*<0.05, \*\**P*<0.01, \*\*\**P*<0.001; ns, not significant, by one-way ANOVA with Dunn’s pairwise method. (**e**) Representative Immunohistochemistry images of NALCN and SLO2.1 in mouse uteri at the indicated time points, with hematoxylin counterstain. SMA, smooth muscle actin. lm, longitudinal muscle; cm, circular muscle; AM, antimesometrial; M, mesometrial. Scale bar, 100µm. We performed NALCN and SLO2.1 immunohistochemistry in tissues from 11 and 8 females, respectively.

**Figure 2.**
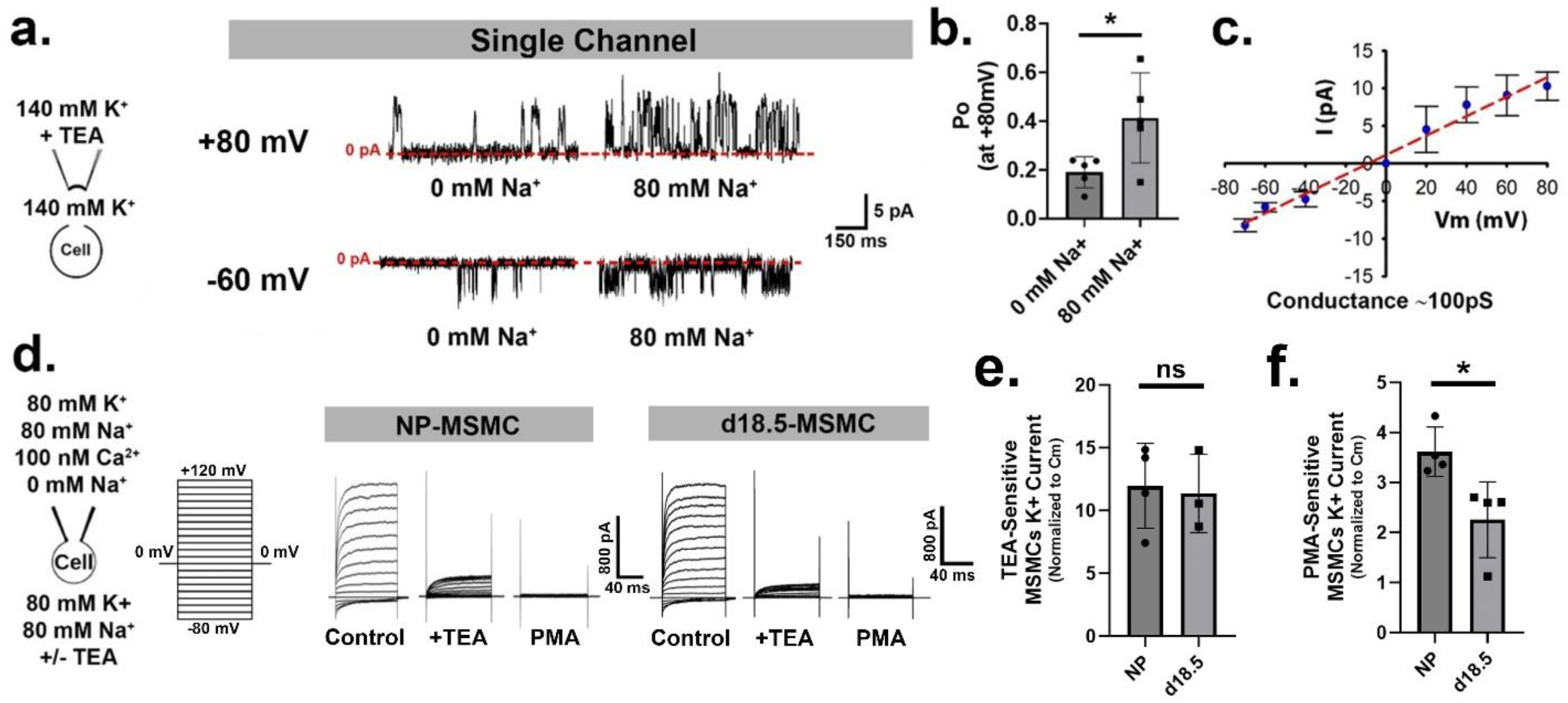
SLO2.1 Sodium-activated potassium currents in mMSMCs. **(a)** Left, schematic of solutions. Right, representative single-channel patch-clamp recordings of SLO2.1 channels from non-pregnant (NP) myometrial smooth muscle cells (MSMCs) at - 60 mV and +80 mV in the presence and absence of 80 mM Na^+^ in a 140 mM symmetrical K^+^ solution. **(b)** Graph showing SLO2.1 channel open probability (Po) at +80 mV in the presence and absence of 80 mM Na^+^. **(c)** Graph showing the mean amplitude and standard deviation (SD) of SLO2.1 single channel current in MSMCs at different voltages (n=4). Channel conductance was calculated from the slope of the fitted line (dotted red line), with a conductance value of 103 pS. **(d)** Left, schematic of whole-cell patch clamp setup, solutions, and voltage steps protocol (Vh = 0 mV, with step pulses from −80 to +120 mV). Right, representative K^+^ currents recorded in MSMCs from NP and day 18.5 (d18.5) females under control conditions, in the presence of the SLO1 (BK) blocker 5 mM TEA (+TEA), and in the presence of the PKC activator and SLO2.1 inhibitor, 1 µM PMA (PMA). **(e)** Graph showing SLO1 (TEA-sensitive) and **(f)** Graph showing SLO2.1 (PMA-sensitive) currents normalized to cell capacitance (Cm) from NP and d18.5 females. Values are (mean ± SD): for **(e)** NP-TEA-Sensitive (11.98 ± 3.37, n=4 cells), and d18.5-TEA-Sensitive (12.38 ± 3.25, n=3 cells). For **(f)** NP-PMA-Sensitive (3.62 ± 0.49, n=4 cells), and d18.5-PMA-Sensitive (2.26 ± 0.76, n=4 cells). \**P* < 0.05; ns, not significant, by unpaired t-test.

## 3. Results

### 3.1. SLO2.1 and NALCN are expressed in mouse myometrial smooth muscle cells

To determine whether SLO2.1 and NALCN are expressed in mouse MSMCs as they are in human MSMCs, we first performed qRT-PCR to examine expression of the related channels *Slo2.1* and *Slo2.2*. We could not detect *Slo2.2* mRNA at any stage but could detect *Slo2.1* mRNA in MSMCs isolated from non-pregnant mice and mice at different stages of pregnancy (Figure 1a, b). *Slo2.1* and *Nalcn* mRNAs were significantly more abundant in uteri of non-pregnant mice and uteri from day post-copulation 7.5 (d7.5) mice than in uteri from mice at term (d18.5) (Figure 1c, d). Immunohistochemistry showed that, in non-pregnant mice, SLO2.1 and NALCN proteins were expressed in the longitudinal and circular smooth muscle layers (Figure 1e, Supplementary Figure 1, and 2). On d7.5, SLO2.1 and NALCN were evident in both mesometrial and antimesometrial myometrial smooth muscle at both implantation and inter-implantation sites. However, NALCN was more abundant in the mesometrial myometrium than in the antimesometrial myometrium. At d14.5 and d18.5, SLO2.1 and NALCN expression in smooth muscle appeared lower than at d7.5. Staining for smooth muscle actin confirmed that SLO2.1 and NALCN were predominantly expressed in smooth muscle. Staining in the decidua was largely non-specific background (Supplementary Figure 1). These findings indicate that SLO2.1 and NALCN are expressed in mouse MSMCs, and that expression decreases late in pregnancy.

### 3.2. Mouse myometrial smooth muscle cells conduct a Na^+^-activated K^+^ current driven by SLO2.1

To assess the presence of Na^+^-activated K^+^ currents in MSMCs isolated from non-pregnant mice, we first conducted single-channel recordings, revealing Na^+^-activated K^+^ channels with an approximate conductance of 100 pS, likely the Na^+^-activated K^+^ channel SLO2.1 (Figure 2 a, b and c). We also detected activity (not shown) of the large-conductance Ca^2+^-activated K^+^ channel SLO1 (BK_Ca_, encoded by *Kcnma1*), which is expressed in MSMCs (Brainard et al., 2009; Ferreira et al., 2019; Khan et al., 1993, 1998; Lorca et al., 2014; Wakle-Prabagaran et al., 2016). We next performed whole-cell patch clamp experiments in which we applied depolarizing voltage clamp steps (10 mV) from −80 mV to +120 mV (Vh= 0 mV) in MSMCs. This revealed a time-dependent, non-inactivating outward K^+^ current (Figure 2d). We suspected that a portion of this current was conducted by SLO1 (Brainard et al., 2009; Ferreira et al., 2019; Khan et al., 1993, 1998; Lorca et al., 2014; Wakle-Prabagaran et al., 2016), so we treated MSMCs from non-pregnant and d18.5 mice with the SLO1 blocker tetraethylammonium (TEA) (5 mM) (Figure 2d, e). These experiments revealed a TEA-sensitive current (60–70% of the total whole-cell current) and a TEA-resistant current (30-40% of the total whole-cell K^+^ current). To confirm that the TEA-resistant current was conducted by SLO2.1, we treated cells with phorbol 12-myristate 13-acetate (PMA). PMA activates protein kinase C, which inhibits SLO2.1, so PMA is an indirect inhibitor of SLO2.1 (Chen et al., 2009; Kaczmarek, 2013; Santi et al., 2006). PMA inhibited the TEA-resistant current in MSMCs from both non-pregnant and d18.5 mice (Figure 2f). We conclude that SLO2.1 channels are responsible for the TEA-resistant K^+^ current in mouse MSMCs. Consistent with our expression data, SLO2.1 current was lower in MSMCs from d18.5 mice than in MSMCs from non-pregnant mice (Figure 2f).

### 3.3. SLO2.1 and NALCN co-localize in non-pregnant mouse MSMCs

SLO2.1 and other Na^+^ channels form complexes in neurons (Hage & Salkoff, 2012; Takahashi & Yoshino, 2015), and SLO2.1 and NALCN form a complex in human MSMCs (Ferreira et al., 2021). To determine whether NALCN co-localizes with SLO2.1 in mouse MSMCs, we conducted *in situ* proximity ligation assays in which we stained mouse MSMCs with antibodies recognizing SLO2.1 and NALCN, along with DNA-tagged secondary antibodies. If the proteins were within 40 nm, DNA ligation and subsequent amplification would occur, leading to fluorescent punctae in the cells. We observed significantly more punctae in cells stained with both anti-NALCN and anti-SLO2.1 antibodies than in cells stained with either antibody alone or with only secondary antibodies (Figure 3). This finding indicates that NALCN and SLO2.1 co-localize in mouse MSMCs.

**Figure 3.**
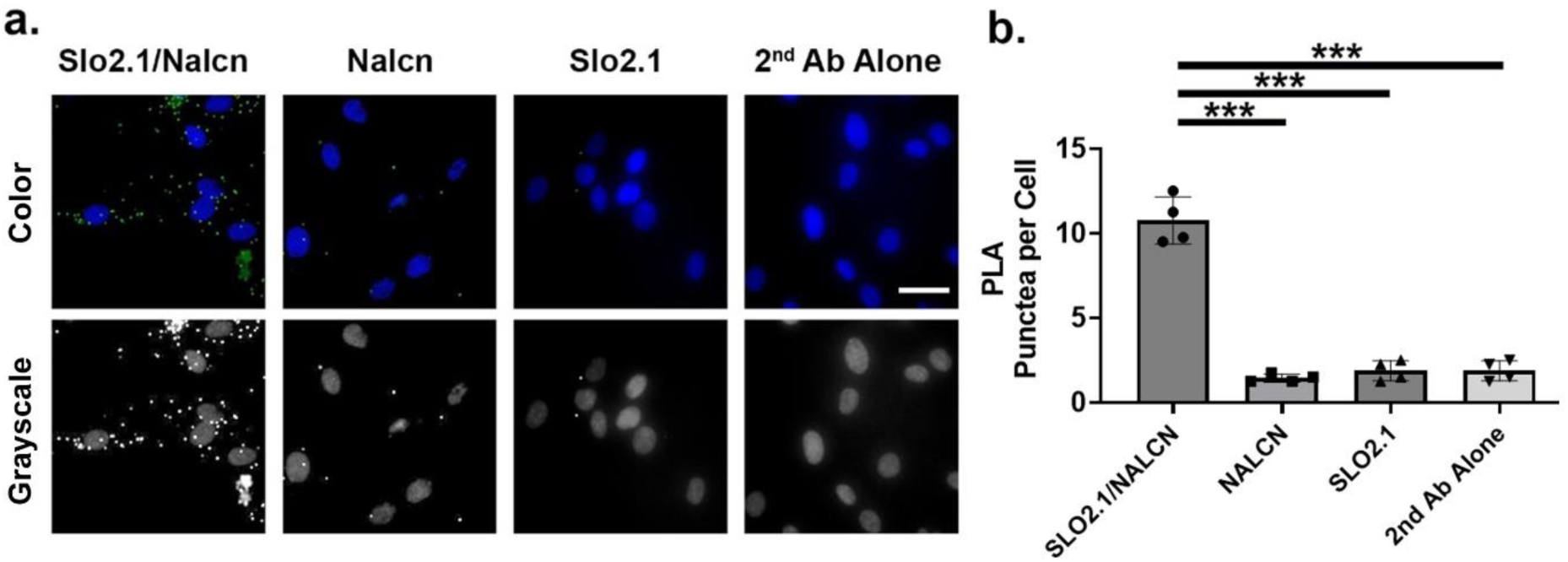
SLO2.1 and NALCN colocalization in mouse MSMCs. (**a**) Representative proximity ligation assay labeling (green) of mouse MSMCs with the indicated single antibodies and antibody combinations. Blue is nuclear stain. Scale bar, 20 µm. (**b**) Average number of punctae in mouse MSMCs with the indicated antibodies. The values are: SLO2.1/NALCN, 10.75 ± 1.40; NALCN, 1.44 ± 0.24; SLO2.1, 1.87 ± 0.59; and 2^nd^ Ab Alone, 1.75 ± 0.95, all from n = 4 (mice). Over 100 cells per condition were counted. Data are presented as mean and standard deviation. \*\*\**P*< 0.001 by one-way ANOVA test with Tukey method.

**Figure 4.**
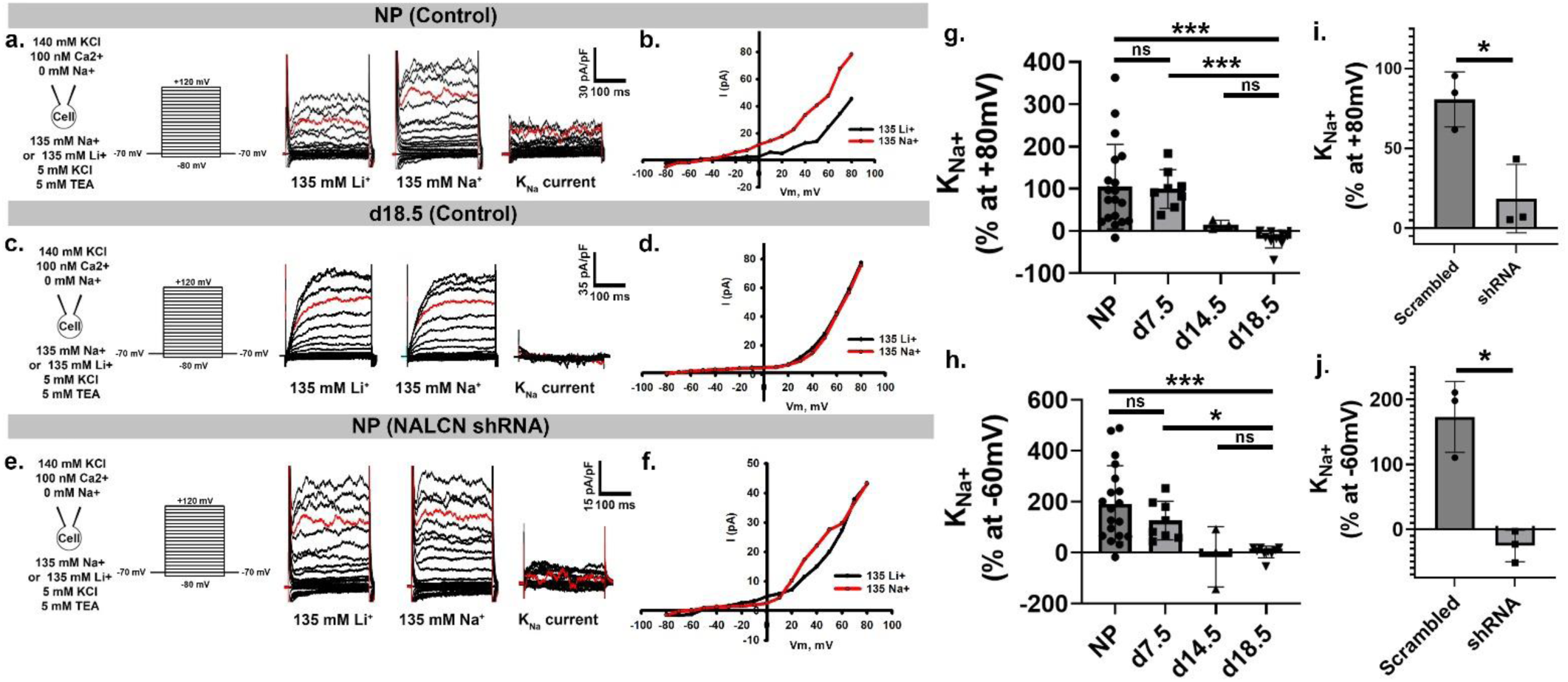
SLO2.1 channel activation by an NALCN-dependent Na^+^ current in mouse MSMCs. (**a, c,** and **e**) Left, schematics of whole-cell bath and pipette solution ionic concentrations and voltage steps applied to the cells. Right, representative whole-cell currents elicited from Vh = −70 mV, and step pulses from −80 to +120 mV, recorded in 135 mM Na^+^ or 135 mM external Li^+^ of MSMCs isolated from **(a)** non-pregnant (NP), **(c)** pregnancy day-18.5 (d18.5), and **(e)** NP mice treated with NALCN-shRNA. The Na^+^-dependent currents were calculated by subtracting traces (135 mM Na^+^ – 135 mM Li^+^). **(b, d,** and **f)** Graphs showing the current-voltage (I-V) relationships for the representative currents: 135 mM Li^+^ or control (black lines) and 135 mM Na^+^ (red lines). **(g** and **h)** Graphs depicting at different stages of pregnancy the percentage of Na^+^-dependent currents at +80 mV and −60 mV, respectively. Data are plotted as mean ± standard deviation. **(g)** Values are: NP, 104.66 ± 99.75, n = 19 cells; d7.5, 99.37 ± 45.79, n = 8 cells; d14.5, 14.84 ± 10.39, n = 3 cells; d18.5, −19.22 ± 20.99, n = 9 cells; **(h)** Values are: NP, 191.44 ± 149.69, n = 19 cells; d7.5, 126.20 ± 76.14, n = 8 cells; d14.5, −16.38 ± 118.5, n = 3 cells; d18.5, 2.13 ± 22.87, n = 9 cells. **(i** and **j)** Graphs depicting in animals treated with Scrambled or NALCN shRNA the percentage of Na^+^-dependent currents at +80 mV and −60 mV, respectively. **(i)** Values are: Scrambled, 80.65 ± 17.25, n = 3 cells, and NALCN shRNA, 18.54 ± 21.41, n = 3 cells. **(j)** Values are: Scrambled, 173.16 ± 54.65, n = 3 cells, and NALCN shRNA, −25.91 ± 24.32, n = 3 cells. \**P* < 0.05, \*\**P* < 0.01, \*\*\**P* < 0.001; ns, not significant, in g and h by one-way ANOVA with Dunn’s pairwise method and in i and j, by paired t-test.

### 3.4. Na^+^ influx through NALCN activates SLO2.1 in MSMCs from non-pregnant and early pregnant mice

We first confirmed that NALCN conducts a Na^+^ current in MSMCs from non-pregnant and d18.5 mice (Supplementary Figure 3). To determine whether Na^+^ influx through NALCN activates SLO2.1 in mouse MSMCs as in human MSMCs (Ferreira et al., 2021), we conducted whole-cell patch clamp experiments on MSMCs isolated from different stages of pregnancy. Substituting 135 mM Li^+^ with 135 mM extracellular Na^+^ led to a 100–200% increase in K^+^ currents at both +80 mV and −60 mV in MSMCs from non-pregnant and early pregnant (d7.5) mice but not in MSMCs from d14.5 and d18.5 (Figure 4a, b, c, d, g, and h). To determine whether the channel responsible for conducting Na^+^ to activate SLO2.1 was NALCN, we conducted similar experiments in non-pregnant mouse MSMCs expressing shRNA specific to NALCN (Figure 4i, and j). In these cells, the Na^+^-activated K^+^ current activated by Na^+^ was 10-20% of that in control MSMCs (Figure 4i, and j). Additionally, Gd^2+^ significantly inhibited this current (Supplementary Figure 4). These findings suggest that, in non-pregnant and early pregnancy stages, the Na^+^ responsible for activating K^+^ efflux through SLO2.1 enters MSMCs through NALCN.

### 3.5. Activation of the NALCN/SLO2.1 complex hyperpolarizes MSMCs from non-pregnant and early pregnancy stage mice

We next wondered whether Na^+^-mediated activation of SLO2.1 and the ensuing K^+^ efflux could lead to membrane hyperpolarization in mouse MSMCs. Thus, we performed flow cytometry with the cationic voltage-sensitive dye, DiSC3(5) (Ferreira et al., 2021). In MSMCs from non-pregnant and d7.5 mice, treatment with 80 mM extracellular Na^+^ led to a respective increase of −28.98 +/− 24.51 and −47.67 +/− 26.02 in DiSC3(5) fluorescence (normalized to Valinomycin), indicating membrane hyperpolarization at these stages (Figure 5b, c, and d). Treatment with 80 mM Li^+^, a NALCN-permeable cation that does not activate SLO2.1, depolarized MSMCs from non-pregnant mice (Figure 5c). Similar experiments with the impermeable cation choline also showed no membrane hyperpolarization, indicating that the Na^+^ effect was not due to osmolarity changes (Ferreira et al. 2021). Conversely, MSMCs from d14.5 and d18.5 mice exhibited a statistically significant depolarization after addition of 80 mM Na^+^ (Figure 5b, d). This indicated that Na^+^ influx at these stages did not activate K^+^ efflux through SLO2.1.

**Figure 5.**
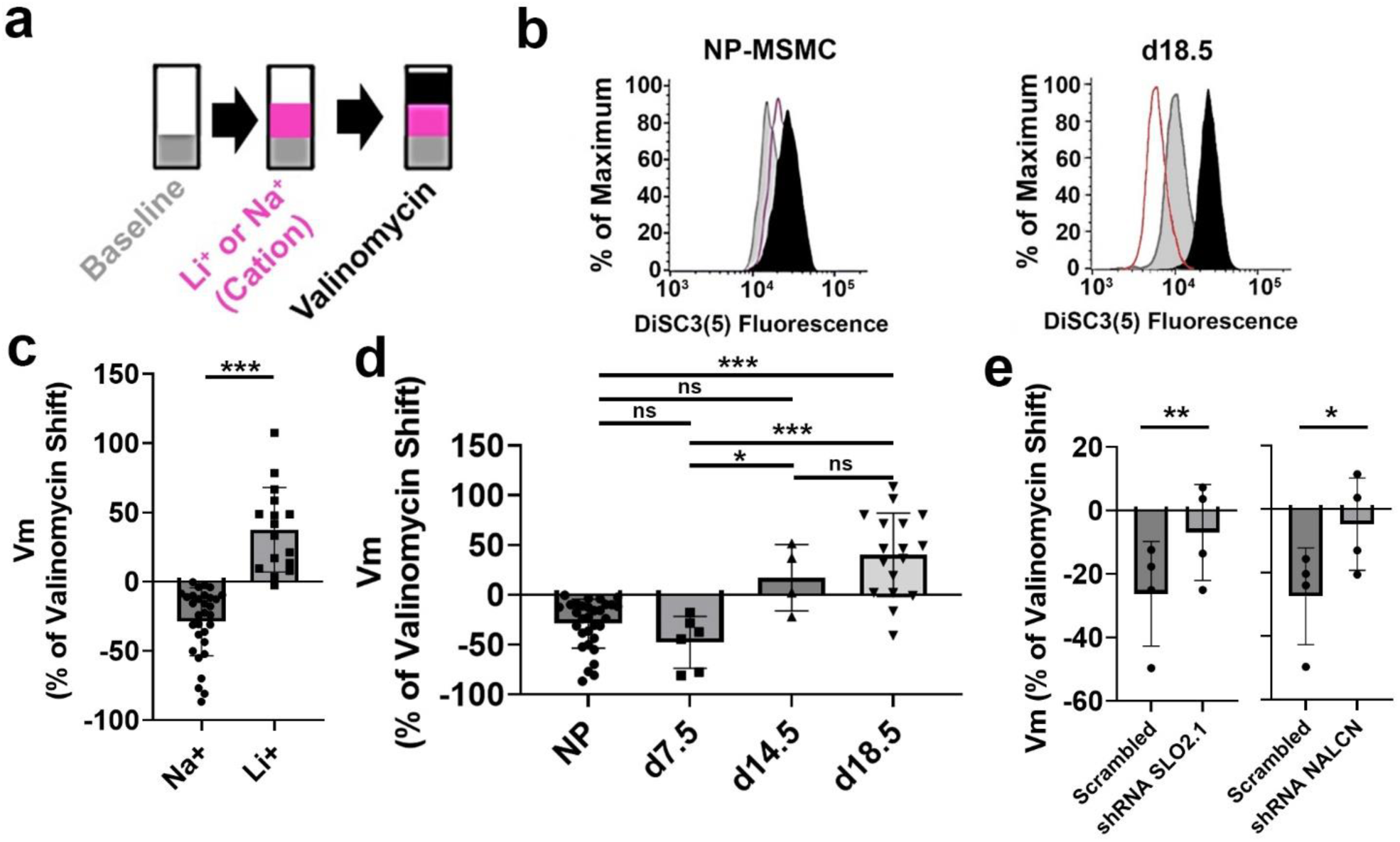
Myometrial smooth muscle cell membrane potential regulation by SLO2.1 and NALCN. (**a**) Experimental scheme. (**b**) Representative images of relative shifts of DiSC3(5) fluorescence induced by 80 mM Na^+^ in MSMCs from non-pregnant (NP) and pregnancy day (d) 18.5 mice. (**c**) Quantification of shifts induced by Na^+^ (−28.22 ± 26.16, n= 27 mice), and Li^+^ (42.83 ± 31.54, n= 13 mice) in MSMCs from NP mice. (**d**) Quantification of shifts induced by Na^+^. Values are NP, −28.98 ± 24.51, n= 31 mice; d7.5, −47.67 ± 26.02, n=6 mice; d14.5, 17.23 ± 33.27, n=4 mice; and d18.5, 40.22 ± 41.78, n=17 mice. (**e, left**) Quantification of shifts induced by Na^+^ in NP mice treated with Scrambled or SLO2.1 shRNA. Values are: Scrambled −26.14 ± 16.47, n=4 mice, and for SLO2.1 shRNA −6.87 ± 15.10, n = 4 mice. (**e, right**) Quantification of shifts induced by Na^+^ in NP mice treated with Scrambled or NALCN shRNA. Values are: Scrambled −27.41 ± 15.22, n=4 mice, and for NALCN shRNA −4.70 ± 14.56, n = 4 mice. (**c to e**) All data were normalized to the fluorescence changes induced by valinomycin and are presented as mean and standard deviation. \*\**P* < 0.01, \*\*\**P* < 0.001, ns, not significant, for **(c)** by unpaired t-test with Mann-Whitney corrections, for **(d)** by one-way ANOVA with Dunn’s pairwise method and in **(e)** by paired t-test.

To confirm that SLO2.1 and NALCN contribute to the Na^+^-induced membrane hyperpolarization, we isolated MSMCs from mice treated with shRNAs specific for *Slo2.1* or *Nalcn*. These MSMCs exhibited significantly smaller Na^+^-dependent membrane hyperpolarization than those that received scrambled shRNA control (Figure 5e). Consistent with these findings, treatment with PMA resulted in Na^+^-dependent depolarization, and treatment with Gd^2+^ or NALCN inhibitors prevented activation of SLO2.1 by Na^+^ influx (Supplementary figure 5). Together, our results suggest that Na^+^ influx through NALCN activates SLO2.1-dependent membrane hyperpolarization in mouse MSMCs.

### 3.6 SLO2.1 and NALCN differentially modulate Ca^2+^ responses in mouse MSMCs across pregnancy

Membrane depolarization in human and mouse MSMCs enhances Ca^2+^ influx via VDCCs (Ferreira et al., 2021), and inhibiting SLO2.1 in human MSMCs leads to increased Ca^2+^ entry through VDCCs (Ferreira et al., 2019, 2021). Thus, we wanted to define the effects of the NALCN/SLO2.1 complex on Ca^2+^ in mouse MSMCs across pregnancy. To do so, we loaded MSMCs with the Ca^2+^ indicator Fluo4-AM. Treatment of MSMCs from non-pregnant mice with 135 mM Li^+^ induced similar Ca^2+^ oscillations (Figure 6a, and c) as those prompted by 80 mM KCl (Supplementary Figure 6a), whereas 135 mM Na^+^ abolished these oscillations (not shown). Treatment of MSMCs from non-pregnant mice in which *Slo2.1* or *Nalcn* were knocked down caused no Ca^2+^ oscillations, indicating that the oscillations depended on the two channels (Figure 6d). Similarly, Li^+^-induced Ca^2+^ oscillations were significantly reduced in cells treated with NALCN inhibitors (Supplementary figure 6). Conversely, treatment with 135 mM Li^+^ did not induce Ca^2+^ oscillations in MSMCs from pregnancy d18.5 mice (Figure 6b, and c). Thus, these results showed that the NALCN/SLO2.1 complex is more active at the beginning than at the end of pregnancy.

**Figure 6.**
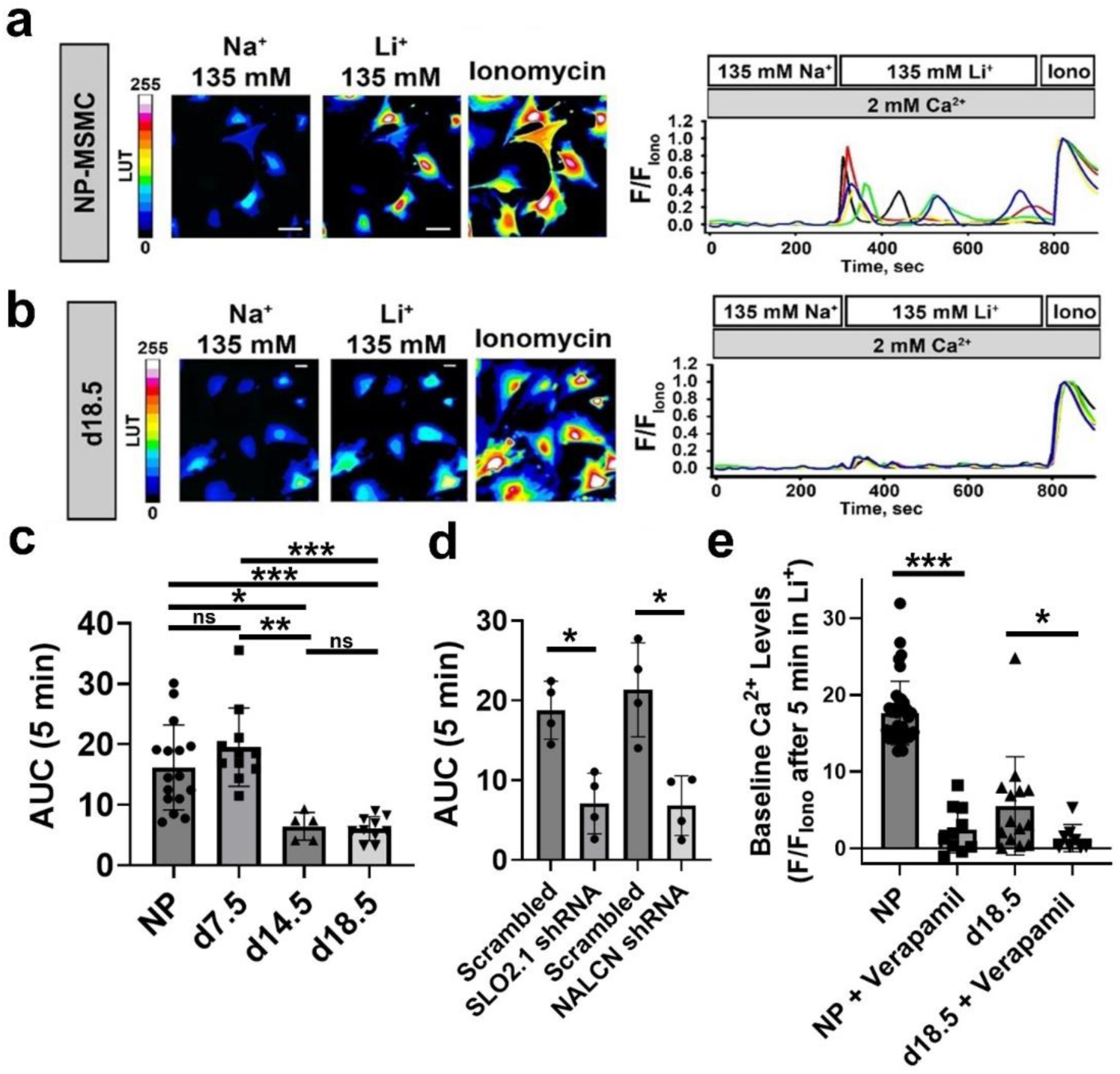
SLO2.1/NALCN regulation of intracellular calcium homeostasis in mouse MSMCs. (**a to b**) Representative fluorescence images (left) and quantitation (right) from MSMCs from non-pregnant (NP) and pregnancy day (d) 18.5 mice loaded with 10 μM Fluo-4 AM in the presence of the indicated ions. (**c**) Graph of the areas under the curve (AUC) of the first 5 min after changing the solutions from 135 mM Na^+^ to 135 mM Li^+^. Values are: NP, 16.17 ± 7.0, n= 16 mice; d7.5, 19.53 ± 6.46, n= 11 mice; d14.5, 6.46 ± 2.29, n= 4 mice; d18.5, 6.01 ± 1.96, n= 10 mice. (**d**) Graph of the areas under the curve (AUC) of the first 5 min after changing the solutions from 135 mM Na^+^ to 135 mM Li^+^ in animals treated with shRNA against SLO2.1, NALCN and paired Scrambled. Values are: SLO2.1 paired scrambled, 18.77 ± 3.62, n= 4 mice; SLO2.1 shRNA, 7.09 ± 3.78, n= 4 mice; NALCN paired scrambled, 21.34 ± 5.88, n= 4 mice; and NALCN shRNA, 6.82 ± 3.74, n= 4 mice. (**e**) Graph of the baseline Ca^2+^ in NP and d18.5 mice, in control conditions and in the presence of the voltage-activated calcium channel inhibitor verapamil. Values are: NP, 17.65 ± 4.15, cells = 37, mice =13; NP + Verapamil, 2.42 ± 3.02, n= 4 mice; d18.5, 6.81 ± 7.93, cells = 10, mice = 4 mice; and d18.5 + Verapamil, 1.33 ± 1.76, cells = 8, mice = 4. All data were normalized to the fluorescence in 2-5 μM ionomycin and 2 mM extracellular Ca^2+^ (Iono), and all data are presented as mean and standard deviation. \**P* < 0.05, ** *P*<0.010, *** *P* < 0.001, and ns, not significant, for **(c)** by one-way ANOVA with Dunn’s pairwise method, for **(d)** by paired t-test, and for **(e)** by unpaired t-test with Mann-Whitney correction.

To confirm that the Ca^2+^ oscillations depended on extracellular Ca^2+^, we repeated the experiment in 0 mM extracellular Ca^2+^ plus 2 mM EGTA. This treatment prevented the Li^+^-induced Ca^2+^ oscillations (Supplementary Figure 6). Furthermore, the proportion of cells responding to Li^+^ was lower when the VDCC inhibitor Verapamil was present (Figure 6e). In summary, our results indicate that Na^+^ influx through NALCN activates SLO2.1, leading to membrane hyperpolarization and reduction of Ca^2+^ oscillations in MSMCs from non-pregnant and early pregnancy stage mice but not in MSMCs from late-pregnancy stage mice.

## 4. Discussion

In previous work, we demonstrated that the Na^+^ channel NALCN and the K^+^ channel SLO2.1 form a functional complex regulating the membrane potential of human MSMCs at the end of pregnancy. Here, we present several lines of evidence indicating that NALCN and SLO2.1 form a functional complex that modulates mouse MSMC membrane potential and excitability throughout all stages of pregnancy. First, we show that SLO2.1 and NALCN mRNA and protein were expressed in mMSMCs throughout pregnancy. Their expression declined in late pregnancy, aligning with the higher MSMC excitability observed at this stage. Second, we show that SLO2.1 currents constitute a significant portion of the total K^+^ current in mouse MSMCs. SLO2.1 currents are reduced in late pregnancy, corroborating our expression data. Third, we demonstrate that NALCN and SLO2.1 are in spatial proximity within mouse MSMC membranes. Fourth, knocking down NALCN resulted in a nearly complete reduction of the of the Na^+^-activated SLO2.1 currents in mouse MSMCs. Fifth, in non-pregnant and early stages of pregnancy, the predominant effect of Na^+^ entry through NALCN is hyperpolarization due to SLO2.1 activation, which results in outward K^+^ currents. This occurs because SLO2.1 is a high-conductance ion channel with much greater conductance than NALCN. Finally, we report that SLO2.1/NALCN complex regulates Ca^2+^ entry through VDCCs in MSMCs from non-pregnant or early pregnancy mice, and its contribution to intracellular calcium regulation is significantly reduced in MSMCs from mice in late pregnancy.

Given our data, we propose the following model (Figure 7): In non-pregnant and early-pregnant mice (quiescent state), Na^+^ influx through NALCN activates SLO2.1 channels, promoting K^+^ efflux and maintaining hyperpolarization of the cell membrane. As a result, voltage-dependent Ca^2+^ channels (VDCCs) remain closed, keeping the uterus in a quiescent state. In contrast, toward the end of pregnancy (active state), both the expression and activity of NALCN and SLO2.1 channels decrease, leading to reduced K^+^ efflux, membrane depolarization, and VDCC activation. This increases intracellular Ca^2+^ and enhances uterine contractility.

**Figure 7.**
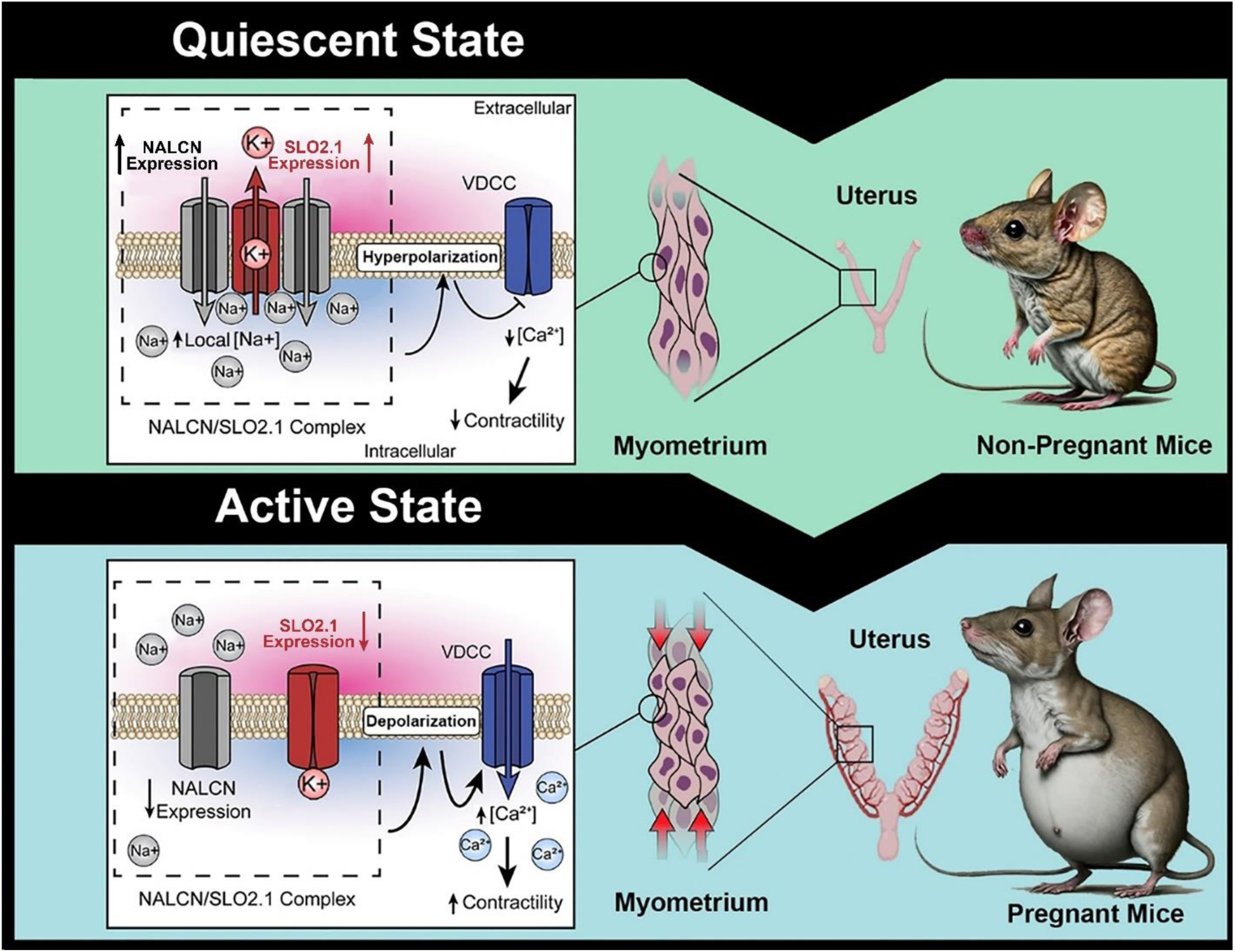
Proposed model by which the SLO2.1/NALCN complex regulates myometrial excitability during pregnancy in mice. In non-pregnant mice and during early pregnancy (“quiescent state”), Na^+^ current through NALCN activates SLO2.1 channels, increasing K^+^ efflux to maintain the cell in a hyperpolarized state. As a result, voltage dependent Ca^2+^ channels (VDCCs) are closed, and the uterus is quiescent. At the end of pregnancy (“active state”), expression of NALCN and SLO2.1 channels and SLO2.1 activity are reduced. The reduced K^+^ efflux depolarizes the membrane, leading to VDCC activation, an increase in intracellular Ca^2+^, and uterine contractility. Image modified from Ferreira et al 2021, https://doi.org/10.1016/j.isci.2021.103210.

Our data are consistent with previous studies showing that human, rat, and mouse MSMCs transition from a hyperpolarized state to a depolarized state at the end of pregnancy (Casteels & Kuriyama, 1965; OSA, 1974; Parkington et al., 1999). This depolarization activates L-type voltage–dependent Ca^2+^ channels, which allow Ca^2+^ influx to both generate an action potential and activate actomyosin contractile machinery (Berridge, 2008). Our mouse model provides insights into the potential role of NALCN in regulating myometrial activity across pregnancy stages. In contrast to previous findings (Reinl et al., 2015b), which suggested a direct effect of NALCN on the muscle inter-burst frequency, our study suggests that the primary role of NALCN is to modulate K^+^ permeability, especially through its interaction with SLO2.1 (Ferreira et al., 2021). In addition to NALCN, other Na^+^ channels such as the TRPC 1, 3 and 6 (Babich et al., 2004; Dalrymple, 2002; Dalrymple et al., 2007; Ku et al., 2006; Lacampagne et al., 1994; Wang et al., 2020; Yang & Sachs, 1989) are expressed in MSMC. However, these channels appear to contribute minimally to Na^+^-activated hyperpolarization in human MSMCs (Ferreira et al., 2021).

In addition to SLO2, other K^+^ channels such as Kir7.1 and SLO1 are expressed in mouse MSMCs (Brainard et al., 2007; Ferreira et al., 2019; Khan et al., 1993; McCloskey et al., 2014). SLO1 channels exhibit high-voltage sensitivity, decreasing their contribution to currents at the resting membrane potential. Kir7.1 channels maintain a hyperpolarized resting membrane potential and low excitability in both mouse and human uterine tissue, peaking in mid-pregnancy in mice (McCloskey et al., 2014). Thus, Kir7.1 could play a role in regulating the hyperpolarized resting membrane potential to maintain uterine quiescence in early pregnancy. However, our study demonstrates that SLO1 and SLO2.1 together contribute to approximately 90% of the K^+^ current in mouse MSMCs, similar to observations in human MSMCs (Ferreira et al., 2019). Further research is needed to fully elucidate the roles of Kir7.1 and TRP channels in mouse MSMCs.

Our data revealed that *Nalcn and SLO2.1* are most highly expressed in MSMCs at d7.5 in pregnancy. At this stage, the myometrium is proliferative, and two recent studies have suggested that NALCN contributes to proliferation. In one paper, NALCN expression in cancer cells, is associated with promoting invasion and metastasis (Folcher et al., 2023). In contrast, Rahrmann et. al, found that loss of NALCN increased the growth of gastric cancer spheroids *in vitro* and cell shedding and metastasis *in vivo* (Rahrmann et al., 2022). More investigation is needed to determine whether NALCN, alone or in combination with SLO2.1, is critical for MSMC proliferation at this early stage of pregnancy. Furthermore, several NALCN accessory subunits, such as Nalf1, have been identified and are thought to be necessary for NALCN function and localization on the plasma membrane (Monteil A, et al., 2023). The role of accessory subunits will be the subject of future investigation.

The results reported in this study should be considered in light of some limitations. First, we did not conduct a priori power analyses to determine sample sizes. Second, with the techniques used in this study, we could not determine whether the reduction in SLO2.1/NALCN functional complex activity at the end of pregnancy was due to decreased channel expression, dissociation of the functional complex, or both. Further research is needed to explore the dynamic interaction between SLO2.1 and NALCN. Finally, there are currently no well-known specific SLO2.1 or NALCN inhibitors or activators that can be considered for therapeutic use. However, we remain vigilant for newly described compounds.

In summary, we demonstrate that the SLO2.1/NALCN functional complex is conserved between mice and humans and functions throughout pregnancy. Future studies should investigate the hormonal modulation of these channels in mice. In human MSMCs, progesterone regulates NALCN expression (Amazu et al., 2020), and oxytocin inhibits SLO2.1 activity (Ferreira et al., 2021). Additional studies should also focus on the interactions with other ion channels and the regulation of NALCN by accessory proteins. Such research could lead to novel drug targets for modulating myometrial contractility.

## Acknowledgments.

We thank Dr. Deborah Frank for critical review of the manuscript. This work was supported by National Institutes of Health grant R01HD088097 (to C.M.S. and S.K.E.), and the Department of Obstetrics and Gynecology at Washington University in St. Louis.

## Author contributions

J.J.F., L.N.K., R.M., A.B, X.M., N.P., C.A., A.Z., G.C.W., S.K.E., and C.M.S. designed, performed, and analyzed the experiments. J.J.F. and L.N.K. prepared figures. J.J.F., L.N.K., S.K.E., and C.M.S. drafted the manuscript. J.J.F., L.N.K., R.M., A.B., X.M., N.P., C.A., A.Z., G.C.W., S.K.E., and C.M.S. edited, revised, and approved the final version of the manuscript.

## Conflict of Interests

Authors declare no conflict of interests.

## Data Availability Statement

The data generated and analyzed during the current study are available from the corresponding author upon reasonable request.

## Notes

### Competing Interest Statement

The authors have declared no competing interest.

### Summary of Updates

The figures and the data has changed since it is going trough the peer reviewed process. This is the latest updated version of the manuscript.

## References.

Amazu, C., Ma, X., Henkes, C., Ferreira, J. J., Santi, C. M., & England, S. K. (2020). Progesterone and estrogen regulate NALCN expression in human myometrial smooth muscle cells. American Journal of Physiology-Endocrinology and Metabolism, 318(4), E441–E452. 10.1152/ajpendo.00320.2019

Anwer, K., Oberti, C., Perez, G. J., Perez-Reyes, N., McDougall, J. K., Monga, M., Sanborn, B. M., Stefani, E., & Toro, L. (1993). Calcium-activated K+ channels as modulators of human myometrial contractile activity. American Journal of Physiology-Cell Physiology, 265(4), C976–C985. 10.1152/ajpcell.1993.265.4.C976

Babich, L. G., Ku, C.-Y., Young, H. W. J., Huang, H., Blackburn, M. R., & Sanborn, B. M. (2004). Expression of Capacitative Calcium TrpC Proteins in Rat Myometrium During Pregnancy1. Biology of Reproduction, 70(4), 919–924. 10.1095/biolreprod.103.023325

Berridge, M. J. (2008). Smooth muscle cell calcium activation mechanisms. The Journal of Physiology, 586(21), 5047–5061. 10.1113/jphysiol.2008.160440

Brainard, A. M., Korovkina, V. P., & England, S. K. (2007). Potassium channels and uterine function. Seminars in Cell & Developmental Biology, 18(3), 332–339. 10.1016/j.semcdb.2007.05.008

Brainard, A. M., Korovkina, V. P., & England, S. K. (2009). Disruption of the maxi-K-caveolin-1 interaction alters current expression in human myometrial cells. Reproductive Biology and Endocrinology, 7(1), 131. 10.1186/1477-7827-7-131

Casteels, R., & Kuriyama, H. (1965). Membrane potential and ionic content in pregnant and non-pregnant rat myometrium. The Journal of Physiology, 177(2), 263–287. 10.1113/jphysiol.1965.sp007591

Chen, H., Kronengold, J., Yan, Y., Gazula, V.-R., Brown, M. R., Ma, L., Ferreira, G., Yang, Y., Bhattacharjee, A., Sigworth, F. J., Salkoff, L., & Kaczmarek, L. K. (2009). The N-Terminal Domain of Slack Determines the Formation and Trafficking of Slick/Slack Heteromeric Sodium-Activated Potassium Channels. The Journal of Neuroscience, 29(17), 5654–5665. 10.1523/JNEUROSCI.5978-08.2009

Dalrymple, A. (2002). Molecular identification and localization of Trp homologues, putative calcium channels, in pregnant human uterus. Molecular Human Reproduction, 8(10), 946–951. 10.1093/molehr/8.10.946

Dalrymple, A., Mahn, K., Poston, L., Songu-Mize, E., & Tribe, R. M. (2007). Mechanical stretch regulates TRPC expression and calcium entry in human myometrial smooth muscle cells. MHR: Basic Science of Reproductive Medicine, 13(3), 171–179*. 10.1093/molehr/gal110

Ferreira, J. J., Amazu, C., Puga-Molina, L. C., Ma, X., England, S. K., & Santi, C. M. (2021). SLO2.1/NALCN a sodium signaling complex that regulates uterine activity. IScience, 24(11), 103210. 10.1016/j.isci.2021.103210

Ferreira, J. J., Butler, A., Stewart, R., Gonzalez-Cota, A. L., Lybaert, P., Amazu, C., Reinl, E. L., Wakle-Prabagaran, M., Salkoff, L., England, S. K., & Santi, C. M. (2019). Oxytocin can regulate myometrial smooth muscle excitability by inhibiting the Na ^+^ -activated K ^+^ channel, Slo2.1. The Journal of Physiology, 597(1), 137–149. 10.1113/JP276806

Folcher, A., Gordienko, D., Iamshanova, O., Bokhobza, A., Shapovalov, G., Kannancheri-Puthooru, D., Mariot, P., Allart, L., Desruelles, E., Spriet, C., Diez, R., Oullier, T., Marionneau-Lambot, S., Brisson, L., Geraci, S., Impheng, H., Lehen’kyi, V., Haustrate, A., Mihalache, A., … Prevarskaya, N. (2023). <scp>NALCN</scp> -mediated sodium influx confers metastatic prostate cancer cell invasiveness. The EMBO Journal, 42(13). 10.15252/embj.2022112198

Hage, T. A., & Salkoff, L. (2012). Sodium-Activated Potassium Channels Are Functionally Coupled to Persistent Sodium Currents. The Journal of Neuroscience, 32(8), 2714–2721. 10.1523/JNEUROSCI.5088-11.2012

Kaczmarek, L. K. (2013). Slack, Slick, and Sodium-Activated Potassium Channels. ISRN Neuroscience, 2013, 1–14. 10.1155/2013/354262

Khan, R. N., Morrison, J. J., Smith, S. K., & Ashford, M. L. J. (1998). Activation of large-conductance potassium channels in pregnant human myometrium by pinacidil. American Journal of Obstetrics and Gynecology, 178(5), 1027–1034. 10.1016/S0002-9378(98)70543-5

Khan, R. N., S K Smith, J J Morrison, & M L Ashford. (1993). Properties of large-conductance K+ channels in human myometrium during pregnancy and labour. Proceedings of the Royal Society of London. Series B: Biological Sciences, 251(1330), 9–15. 10.1098/rspb.1993.0002

Knock, G. A., Smirnov, S. V., & Aaronson, P. I. (1999). Voltage-gated K ^+^ currents in freshly isolated myocytes of the pregnant human myometrium. The Journal of Physiology, 518(3), 769–781. 10.1111/j.1469-7793.1999.0769p.x

Ku, C. Y., Babich, L., Word, R. A., Zhong, M., Ulloa, A., Monga, M., & Sanborn, B. M. (2006). Expression of Transient Receptor Channel Proteins in Human Fundal Myometrium in Pregnancy. Journal of the Society for Gynecologic Investigation, 13(3), 217–225. 10.1016/j.jsgi.2005.12.007

Lacampagne, A., Gannier, F., Argibay, J., Garnier, D., & Le Guennec, J.-Y. (1994). The stretch-activated ion channel blocker gadolinium also blocks L-type calcium channels in isolated ventricular myocytes of the guinea-pig. Biochimica et Biophysica Acta (BBA) - Biomembranes, 1191(1), 205–208. 10.1016/0005-2736(94)90250-X

Lorca, R. A., Prabagaran, M., & England, S. K. (2014). Functional insights into modulation of BKCa channel activity to alter myometrial contractility. Frontiers in Physiology, 5. 10.3389/fphys.2014.00289

Monteil A, Guérineau NC, Gil-Nagel A, Parra-Diaz P, Lory P, Senatore A. (2023). New insights into the physiology and pathophysiology of the atypical sodium leak channel NALCN. Physiol Rev. 2024 Jan 1;104(1):399–472. doi: 10.1152/physrev.00014.2022. PMID: 37615954.

McCloskey, C., Rada, C., Bailey, E., McCavera, S., van den Berg, H. A., Atia, J., Rand, D. A., Shmygol, A., Chan, Y., Quenby, S., Brosens, J. J., Vatish, M., Zhang, J., Denton, J. S., Taggart, M. J., Kettleborough, C., Tickle, D., Jerman, J., Wright, P., … Blanks, A. M. (2014). The inwardly rectifying K ^+^ channel <scp>KIR</scp> 7.1 controls uterine excitability throughout pregnancy. EMBO Molecular Medicine, 6(9), 1161–1174. 10.15252/emmm.201403944

Osa, T. (1974). AN INTERACTION BETWEEN THE ELECTRICAL ACTIVITIES OF LONGITUDINAL AND CIRCULAR SMOOTH MUSCLES OF PREGNANT MOUSE UTERUS. The Japanese Journal of Physiology, 24(2), 189–203. 10.2170/jjphysiol.24.189

Parkington, H. C., Tonta, M. A., Brennecke, S. P., & Coleman, H. A. (1999). Contractile activity, membrane potential, and cytoplasmic calcium in human uterine smooth muscle in the third trimester of pregnancy and during labor. American Journal of Obstetrics and Gynecology, 181(6), 1445–1451. 10.1016/S0002-9378(99)70390-X

Rahrmann, E. P., Shorthouse, D., Jassim, A., Hu, L. P., Ortiz, M., Mahler-Araujo, B., Vogel, P., Paez-Ribes, M., Fatemi, A., Hannon, G. J., Iyer, R., Blundon, J. A., Lourenço, F. C., Kay, J., Nazarian, R. M., Hall, B. A., Zakharenko, S. S., Winton, D. J., Zhu, L., & Gilbertson, R. J. (2022). The NALCN channel regulates metastasis and nonmalignant cell dissemination. Nature Genetics, 54(12), 1827–1838. 10.1038/s41588-022-01182-0

Reinl, E. L., Cabeza, R., Gregory, I. A., Cahill, A. G., & England, S. K. (2015a). Sodium leak channel, non-selective contributes to the leak current in human myometrial smooth muscle cells from pregnant women. Molecular Human Reproduction, 21(10), 816–824. 10.1093/molehr/gav038

Reinl, E. L., Cabeza, R., Gregory, I. A., Cahill, A. G., & England, S. K. (2015b). Sodium leak channel, non-selective contributes to the leak current in human myometrial smooth muscle cells from pregnant women. Molecular Human Reproduction, 21(10), 816–824. 10.1093/molehr/gav038

Reinl, E. L., Zhao, P., Wu, W., Ma, X., Amazu, C., Bok, R., Hurt, K. J., Wang, Y., & England, S. K. (2018). Na+-Leak Channel, Non-Selective (NALCN) Regulates Myometrial Excitability and Facilitates Successful Parturition. Cellular Physiology and Biochemistry, 48(2), 503–515. 10.1159/000491805

Santi, C. M., Ferreira, G., Yang, B., Gazula, V.-R., Butler, A., Wei, A., Kaczmarek, L. K., & Salkoff, L. (2006). Opposite Regulation of Slick and Slack K ^+^ Channels by Neuromodulators. The Journal of Neuroscience, 26(19), 5059–5068. 10.1523/JNEUROSCI.3372-05.2006

Takahashi, I., & Yoshino, M. (2015). Functional coupling between sodium-activated potassium channels and voltage-dependent persistent sodium currents in cricket Kenyon cells. Journal of Neurophysiology, 114(4), 2450–2459. 10.1152/jn.00087.2015

Wakle-Prabagaran, M., Lorca, R. A., Ma, X., Stamnes, S. J., Amazu, C., Hsiao, J. J., Karch, C. M., Hyrc, K. L., Wright, M. E., & England, S. K. (2016). BK _Ca_ channel regulates calcium oscillations induced by alpha-2-macroglobulin in human myometrial smooth muscle cells. Proceedings of the National Academy of Sciences, 113(16). 10.1073/pnas.1516863113

Wang, H., Cheng, X., Tian, J., Xiao, Y., Tian, T., Xu, F., Hong, X., & Zhu, M. X. (2020). TRPC channels: Structure, function, regulation and recent advances in small molecular probes. Pharmacology & Therapeutics, 209, 107497. 10.1016/j.pharmthera.2020.107497

Yang, X.-C., & Sachs, F. (1989). Block of Stretch-Activated Ion Channels in *Xenopus* Oocytes by Gadolinium and Calcium Ions. Science, 243(4894), 1068–1071. 10.1126/science.2466333

