## Supplementary Data for "SLO2.1/NALCN Functional Complex Activity in Mouse Myometrial Smooth Muscle Cells During Pregnancy"

**Supplementary Figures.**

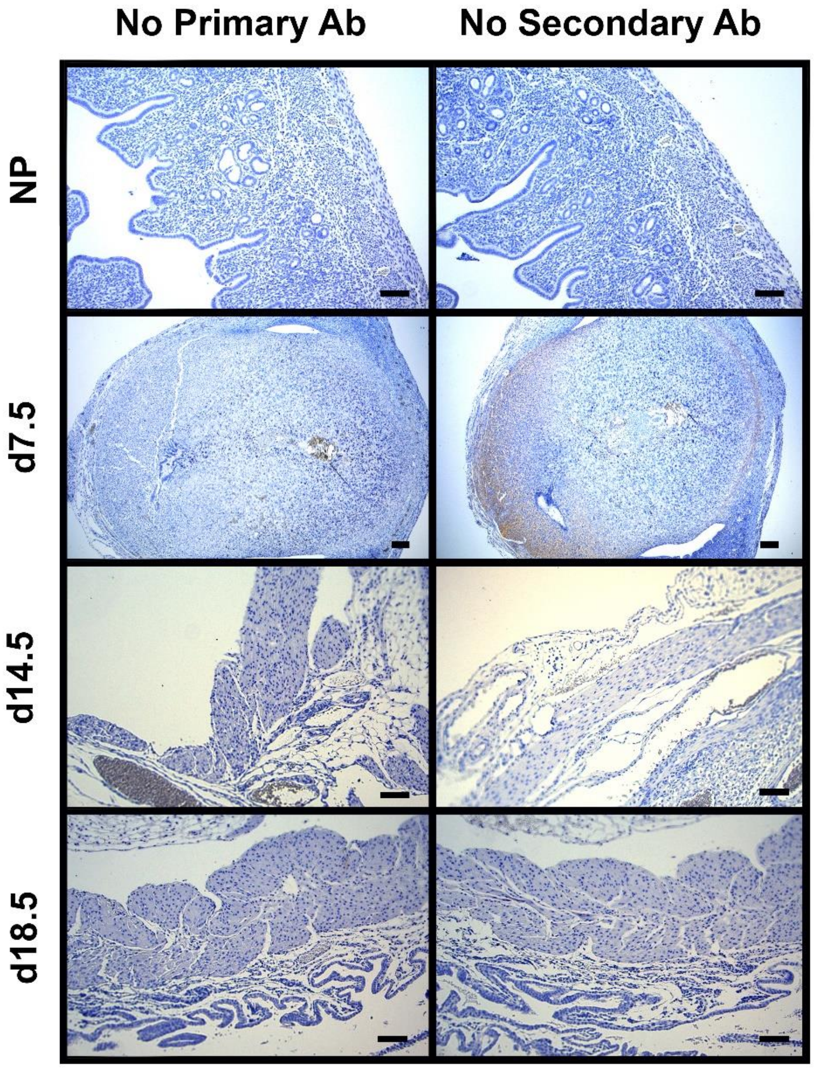


**Supplementary Figure 1. SLO2.1 and NALCN expression in uterus and myometrial smooth muscle cells.** Immunohistochemistry control with No Primary Antibody and No Secondary antibody in mouse uteri at the indicated time points, with hematoxylin counterstain. Scale bar, 100μm.

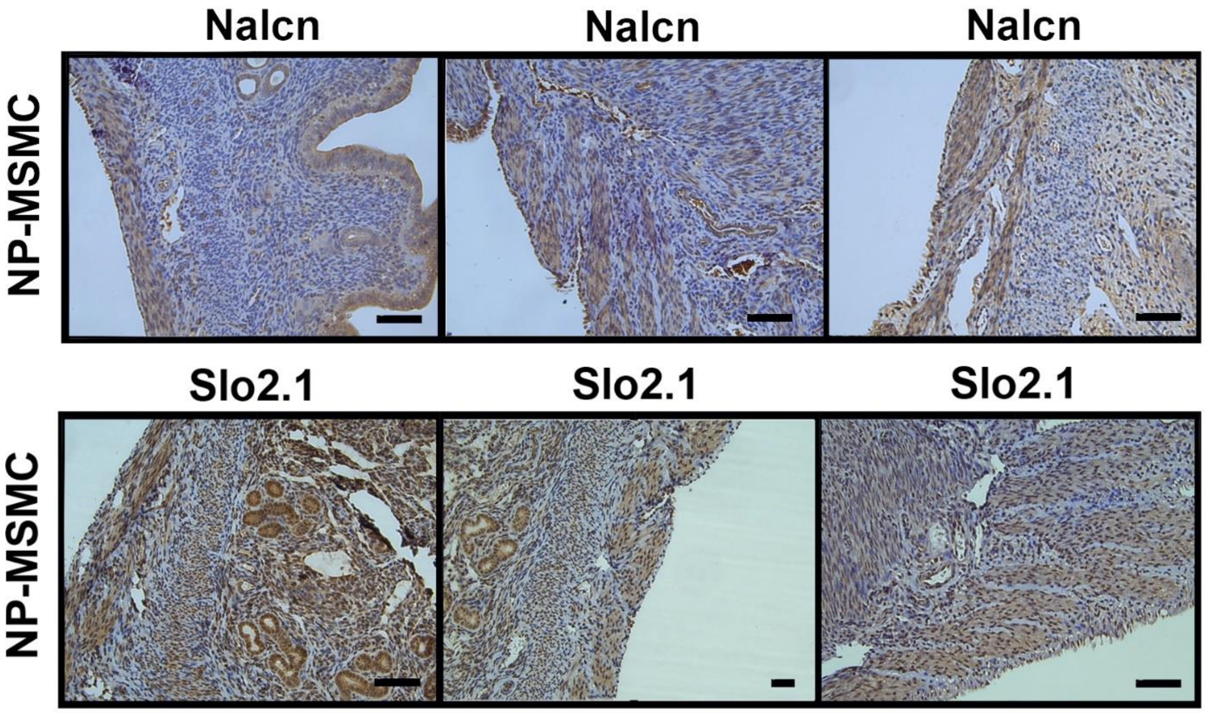


**Supplementary Figure 2. Representative images of SLO2.1 and NALCN expression in uterus and myometrial smooth muscle cells.** Immunohistochemistry of NALCN (n=3) and SLO2.1 (n=3) in mouse uteri, with hematoxylin counterstain. Scale bar, 100μm. We performed immunohistochemistry in 11 females with NALCN antibody and 8 females with SLO2.1 antibody.

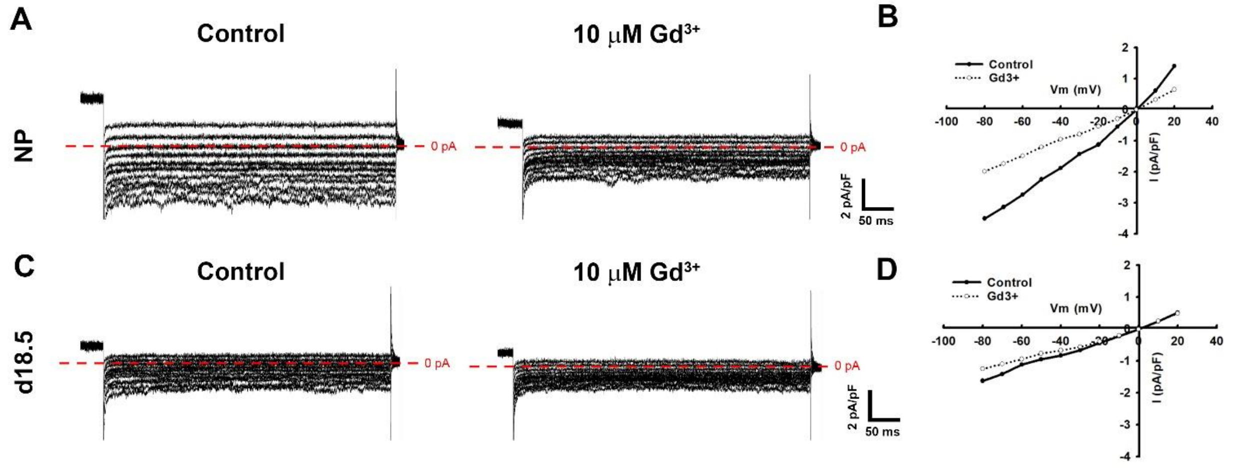


**Supplementary Figure 3.** Representative traces of NALCN and $Gd^{3+}$ sensitive currents in freshly isolated mMSMCs from Non-Pregnant (NP) and Pregnancy day 18.5 (d18.5). **(A and C)** Representative traces of leak current evoked from NP mMSMC **(A)** and d18.5 MSMC **(C)** by using a voltage step protocol with 5 mV increments from -80 mV to +20 mV, before and after treatment with 10 μM $Gd^{3+}$. **(B and D)** Current-voltage relationships obtained from A and C before (black circles) and after (white circles) treatment with 10 μM $Gd^{3+}$. Cells were held at 0 mV and currents were elicited by stepping from -55 mV (50 ms) to +40 mV for 100 ms (to inactivate voltage-gated $Ca^{2+}$ and $K_+$ channels) followed by 5 mV steps (500 ms) from -80 mV to +20 mV using the pClamp 10.6 software (Molecular Devices). Currents were normalized to cell capacitance.

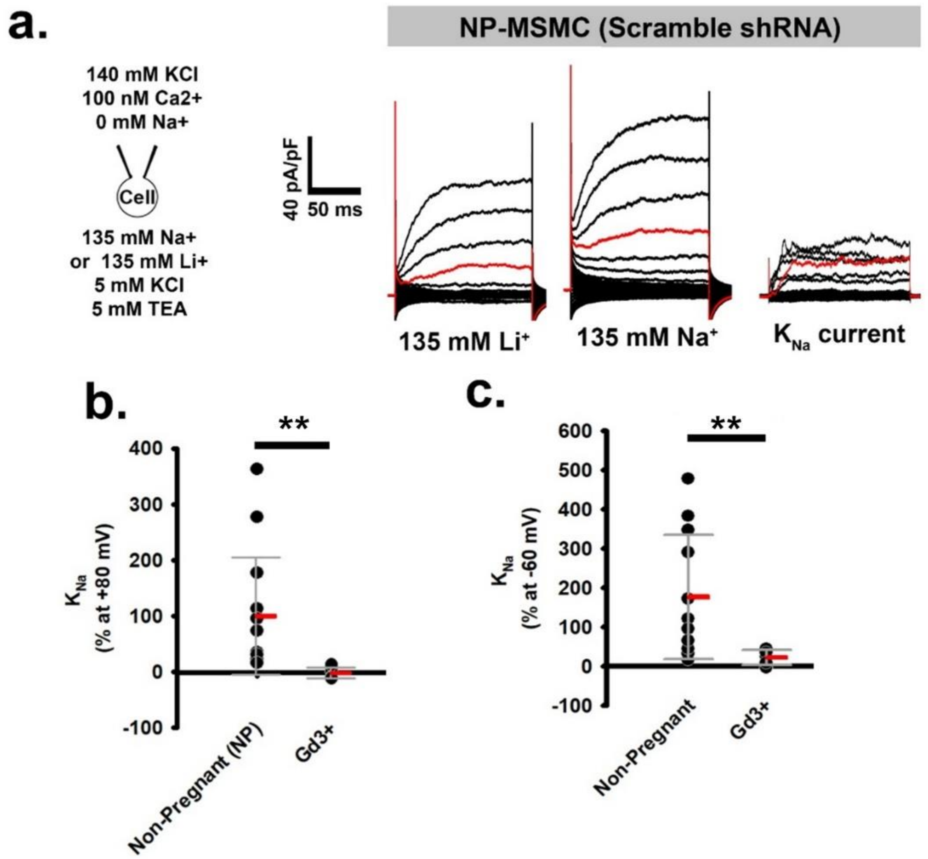


**Supplementary Figure 4. SLO2.1 channel activation by an NALCN-dependent $Na^+$ current in mouse MSMCs.** (**a**) Left, schematics of whole-cell bath and pipette ionic concentrations. Right, representative whole-cell currents elicited from Vh = -70 mV, and step pulses from -80 to +120 mV, recorded in 135 mM $Na^+$ or 135 mM external $Li^+$ of MSMCs isolated from non-pregnant NP mice treated with scramble-shRNA. The $Na^+$-dependent currents were calculated by subtracting traces (135 mM $Na^+$ – 135 mM $Li^+$). (**b** and **c**) Graphs depicting the percentage of $Na^+$-dependent currents at +80 mV and -60 mV, respectively. Data are plotted as the mean and standard deviation. (**b**) Values are NP, 93.50 ± 105.39, n= 10; and $Gd^{3+}$, -0.11±10.14, n=5. (**c**) Values are NP, 173.63 ± 161.17, n= 10; and $Gd^{3+}$, 22.49±19.0, n=5. $^{**}P < 0.01$ by unpaired t-test with or without Mann-Whitney corrections.

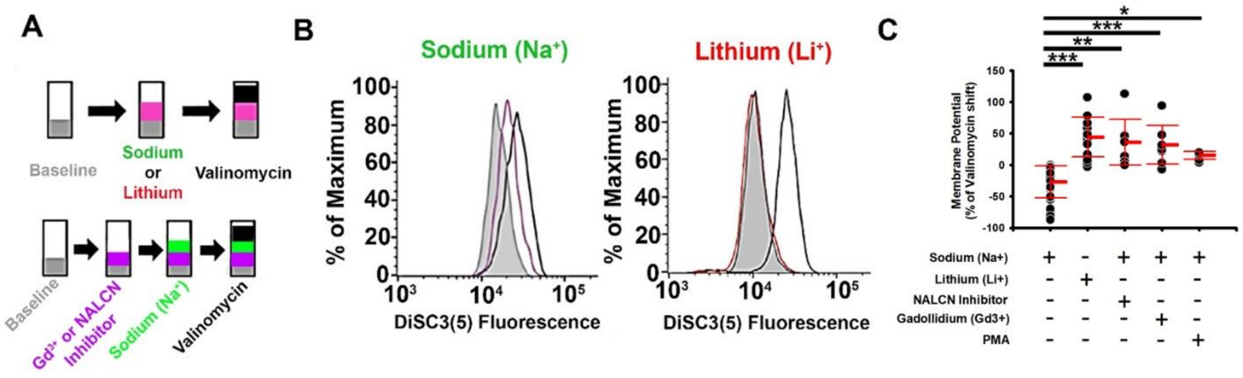


**Supplementary Figure 5. Myometrial smooth muscle cell membrane potential regulation by SLO2.1 and NALCN.** (**a**) Experimental scheme. (**b**) Representative images of relative shifts of DiSC3(5) fluorescence induced by 80 mM $Na^+$ and $Li^+$ in MSMCs from non-pregnant (NP). (**c**) Quantification of shifts induced by $Na^+$ (-28.36 ± 25.24, n= 29), $Li^+$ (42.83 ± 31.54, n= 13), 80 mM $Na^+$ + 50 μM CP96345 (NALCN Inhibitor) (Ferreira et. al. 2021) (34.64 ± 36.52, n=8), 80 mM $Na^+$ + 10 μM $Gd^{3+}$ (30.68 ± 30.74, n=8), and 1 μM phorbol 12-myristate 13-acetate (PMA) (13.82 ± 6.4, n=6). All data were normalized to the fluorescence changes induced by valinomycin and are presented as mean and standard deviation. ***P* < 0.01, ****P* < 0.001 by one-way ANOVA test with Tukey method.

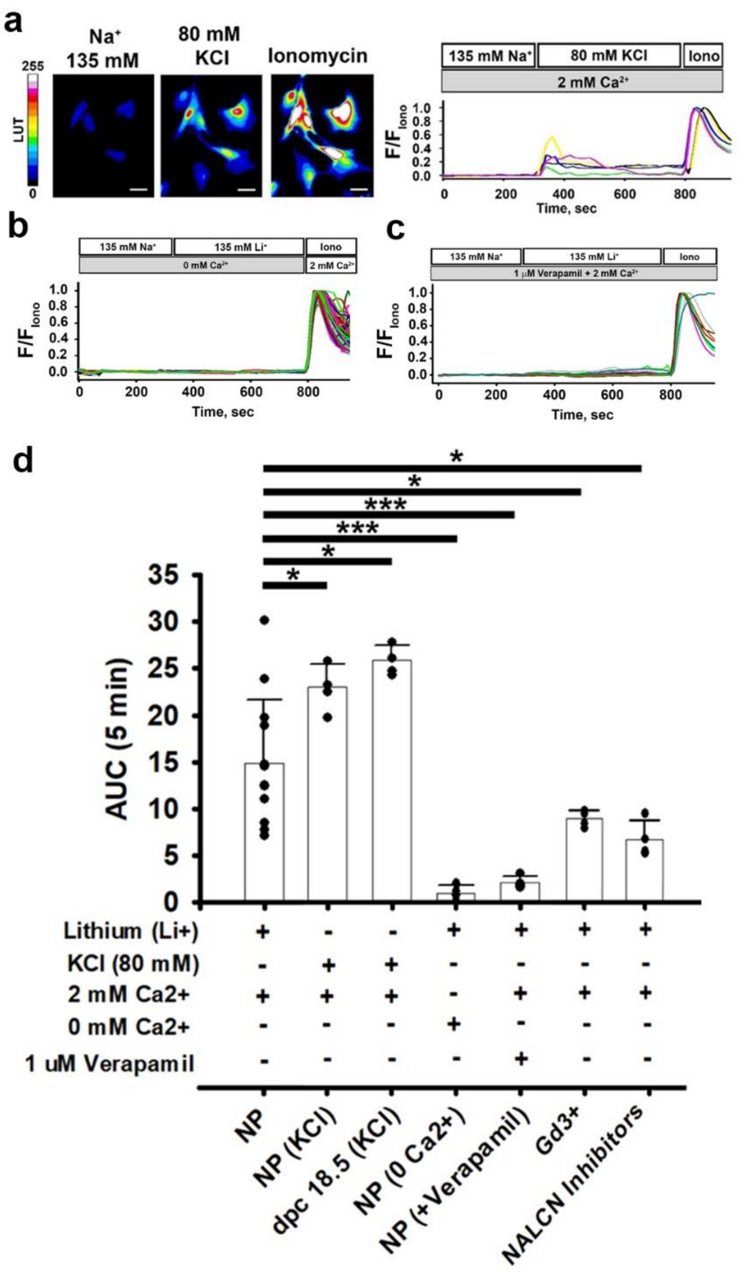


**Supplementary Figure 6. SLO2.1/NALCN regulation of intracellular calcium homeostasis in mouse MSMCs.** (**a, left**) Representative fluorescence images, (**a right, b, and c**) Representative fluorescence traces from non-pregnant (NP) loaded with 10 μM Fluo-4 AM in the presence of extracellular calcium and KCl (a), in absence of extracellular calcium and $Li^+$ (b) and in the presence of Calcium, $Li^+$ and the VDCC inhibitor Verapamil (c). (**d**) Graph of the areas under the curve (AUC) of the first 5 min after changing the solutions from 135 mM $Na^+$ to 135 mM $Li^+$, or from 4 mM KCl to 80 mM KCl in different conditions. All data were normalized to the fluorescence in 2-5 μM ionomycin and 2 mM extracellular $Ca^{2+}$ (Iono), and all data are presented as mean and standard deviation. *$P < 0.05$, and *** $P < 0.001$ by unpaired t-test with or without Mann-Whitney corrections.
